# Are trade-offs between flexibility and efficiency in systematic conservation planning avoidable ?

**DOI:** 10.1101/775072

**Authors:** Sabrine Drira, Frida Ben Rais Lasram, Tarek Hattab, Yunne Jai Shin, Amel Ben Rejeb Jenhani, François Guilhaumon

## Abstract

Species distribution models (SDMs) have been proposed as a way to provide robust inference about species-specific sites suitabilities, and have been increasingly used in systematic conservation planning (SCP) applications. However, despite the fact that the use of SDMs in SCP may raise some potential issues, conservation studies have overlooked to assess the implications of SDMs uncertainties. The integration of these uncertainties in conservation solutions requires the development of a reserve-selection approach based on a suitable optimization algorithm. A large body of research has shown that exact optimization algorithms give very precise control over the gap to optimality of conservation solutions. However, their major shortcoming is that they generate a single binary and indivisible solution. Therefore, they provide no flexibility in the implementation of conservation solutions by stakeholders. On the other hand, heuristic decision-support systems provide large amounts of sub-optimal solutions, and therefore more flexibility. This flexibility arises from the availability of many alternative and sub-optimal conservation solutions. The two principles of efficiency and flexibility are implicitly linked in conservation applications, with the most mathematically efficient solutions being inflexible and the flexible solutions provided by heuristics suffering sub-optimality. In order to avoid the trade-offs between flexibility and efficiency in systematic conservation planning, we propose in this paper a new reserve-selection framework based on mathematical programming optimization combined with a post-selection of SDM outputs. This approach leads to a reserve-selection framework that might provide flexibility while simultaneously addressing efficiency and representativeness of conservation solutions and the adequacy of conservation targets. To exemplify the approach we a nalyzed an experimental design crossing pre- and post-selection of SDM outputs versus heuristics and exact mathematical optimizations. We used the Mediterranean Sea as a biogeographical template for our analyses, integrating the outputs of 8 SDM techniques for 438 fishes species.

## I Introduction

In response to declining wildlife populations and concerns about the anthropogenic footprint on biodiversity (Cardinale et al., 2012; Worm et al., 2006), Systematic Conservation Planning (SCP) has rapidly emerged as a viable approach for identifying protected area systems (set of sites were human activities are restricted or forbiden, hereafter called conservation solutions) that will efficiently meet clearly and transparently defined objectives for biodiversity conservation (Margules and Pressey, 2000; Possingham et al., 2006). SCP is build to satisfy together a set of principles relevant to both the setting of conservation objectives, the identification of conservation solutions and their implementation in conservation actions on the ground (Possingham et al., 2006). Among these principles, adequacy, efficiency and flexibility are key ones (Amorim et al., 2014; Possingham et al., 2006).

An adequate conservation solution grants the persistence of the biodiversity features contained within it (Possingham et al., 2006). In conservation applications, the principle of adequacy is generally addressed by setting quantitative conservation targets that ensure the persistence of all biodiversity features occurring in a region (often species; (Amorim et al., 2014; Carvalho et al., 2016; Giakoumi et al., 2016), but can be functional types (Magris et al., 2017) or habitats (Davies et al., 2017; Drira et al., 2019)). Ideally, conservation targets should relate directly to the probability of species persistence (Araújo and Williams, 2000). As these kinds of data are usually unavailable or incomplete (Araújo et al., 2002), a widely used indirect approach is to set different targets for species with different range sizes, such that species with restricted ranges (who tend to present lower local abundance and higher demographic stochasticity, thus higher extinction risk) have more ambitious targets in terms fo the proportion of their range that should be emcopassed in the conservation solution than widespread species (Harnik et al., 2012; Rodrigues et al., 2004). However, even for such an indirect approach, accurate and complete information about species’ spatial occurrence is required if one wants to set targets granting adequacy on the ground. Biases in species distribution data, known as the Wallacean shortfall, are legion (Bini et al., 2006; Lomolino, 2004; Terribile et al., 2012; Whittaker et al., 2005). This is especially worrisome when trying to plan conservation strategies (Bini et al., 2006; Kadej et al., 2015). Indeed, the selection of protected areas is directly affected by the rates of both omission (false species absences) and commission (false species presences) errors, and using inconsistent distribution data lead to overstated conservation efforts towards well-surveyed areas, whereas unsurveyed areas are disregarded in spite of their potential importance (Rodrigues et al., 2004; Rondinini et al., 2006). To overcome these bias in data, species distribution models (SDMs, also called ecological niche models) have been proposed as a way to provide robust inference about species-specific sites suitabilities. These models use various correlative statistical methods to associate the observed species occurrence patterns within surveyed areas with environmental predictor variables and infer species distribution over all the study area (Elith and Leathwick, 2009). As the different statistical methods used to build SDMs are differently sensitive to geographical range properties (Marmion et al., 2009), including species prevalence (Wisz et al., 2008), they yield variable predictions about species distributions (e.g. Elith et al., 2006; Guisan et al., 2007; Roura-Pascual et al., 2009), generating substantial differences in terms of conservation solutions (Carvalho et al., 2011; Lentini and Wintle, 2015; Loiselle et al., 2003). In this context, the use of SDMs in SCP may raise some potential issues as our ability to evaluate the adequacy of conservation solutions depends strongly on the variability and the quality of SDM outputs (Haight et al., 2000; Rondinini et al., 2006; Wilson et al., 2009). An efficient reserve network is one that is adequate at the least possible cost (or smallest area included in the conservation solution (Margules and Pressey, 2000; Sarkar, 2006). Improving the efficiency of conservation solutions is imperative in a current context of limited conservation budgets (Carvalho et al., 2011; Possingham et al., 2006; Sarkar, 2006). The achievement of efficiency strongly depends on the particular reserve-selection algorithm used to solve the conservation problem, or on its parameterization. A large body of research has shown that exact optimization algorithms (implemented via. mathematical programming) give very precise control over the gap to optimality of conservation solutions (i.e. the degree of solution efficiency), allowing to avoid the waste of scarce conservation resources (Camm et al., 1996; Önal, 2004; Underhill, 1994). However, a major shortcoming of exact reserve-selection algorithms is that they generate a single binary and indivisible solution. They yield no information regarding both the relative importance of selected sites and the opportunity that sites outside the optimal conservation solution represent for the achievement of conservation targets. Therefore, they provide no flexibility in the implementation of conservation solutions by stakeholders (Wilhere et al., 2008). Yet, flexibility, the opportunity to choose alternative sites, is central to SCP, empowering stakeholders to schedule conservation actions and negotiate the inclusion of sites having particular ecological, social, or political interests (Wilson et al., 2009). In the pursuit of flexibility, heuristic decision-support systems (implemented via random search algorithms (e.g. Marxan, Ball et al., 2009) or greedy algorithms (e.g. Zonation, Moilanen and Ball, 2009)), providing large amounts of sub-optimal solutions (containing more sites than necessary or capturing less conservation features than feasible with the same amount of resources), have been widely supported (Vanderkam et al., 2007). Indeed, heuristics allow quantifying the importance of all sites to achieve the conservation goals and therefore the possibility of replacing them. This is based on the calculation of an index, called irreplaceability or optimacity, expressed as the selection frequency of each planning unit across the set of sub-optimal solutions (Wilhere et al., 2008). Hence, the flexibility associated with heuristics-based conservation solutions arises from the availability of many alternative and sub-optimal conservation solutions. As such, in the current context, the two principles of efficiency and flexibility are implicitly linked in conservation applications, with the most mathematically efficient solutions being inflexible and the flexible solutions provided by heuristics suffering sub-optimality. By screening the literature for conservation applications that used SDMs predictions as inputs for SCP we picture the different strategies used to deal with both SDM uncertainties and reserve-selection efficiency (Appendix 1). We have identified four possible approaches resulting from the combinations of two strategies for the treatment of SDM uncertainties (i.e. a pre-selection vs a post-selection approach, see Meller et al., 2014) and two methods used for reserve selection (i.e. heuristics vs mathematical programming) (Figure 1). Today, the use of SDMs is becoming increasingly advocated (Bailey and Thompson, 2009; Elith and Leathwick, 2009), and more frequent in SCP applications (Appendix 1). In the main, conservation studies have overlooked to assess the implications of SDM uncertainties and have focused on a single and a priori-chosen statistical method (Appendix 1, Bailey and Thompson, 2009). Others studies have considered a wide range of models to predict species distributions and used only the best model identified in terms of predictive performance (e.g. (Leach et al., 2013; Passoni et al., 2017; Walther and Pirsig, 2017)). This risks the possibility that a conservation solution based on one particular model might not meet the required target identified under a different good model (Zhang et al., 2015). Proceeding further, some studies used ensemble approaches to summarize the information across multiple model predictions, this takes the uncertainty between different statistical methods into account by yielding predictions representing a central tendency across model predictions (e.g. using median or averaged predicted values) (e.g. (Alagador et al., 2016; Bush et al., 2014; Faleiro and Loyola, 2013). Building protected area systems using ensemble species distribution predictions (called hereafter the “pre-selection approach”) masks the variability in conservation outcomes induced by different SDMs (Meller et al., 2014). To overcome this limitation, Meller et al. (2014) introduced a “post-selection” approach where different distribution scenarios are constructed by randomly sampling across the full range of model predictions and used as inputs for SCP algorithms (Figure 1). This latter approach provides more reliable conservation outcomes, notably showing a better representation of rare species than using the pre-selection approach (Meller et al., 2014). Reserve-selection efficiency has been less considered when implying SDM in the SCP process. Most conservation applications used heuristic algorithms. Only 4% of the reviewed publications have implemented mathematical programming to provide optimal conservation solutions, all of these three studies used a pre-selection approach (see review in Appendix 1).

**Figure 1:**
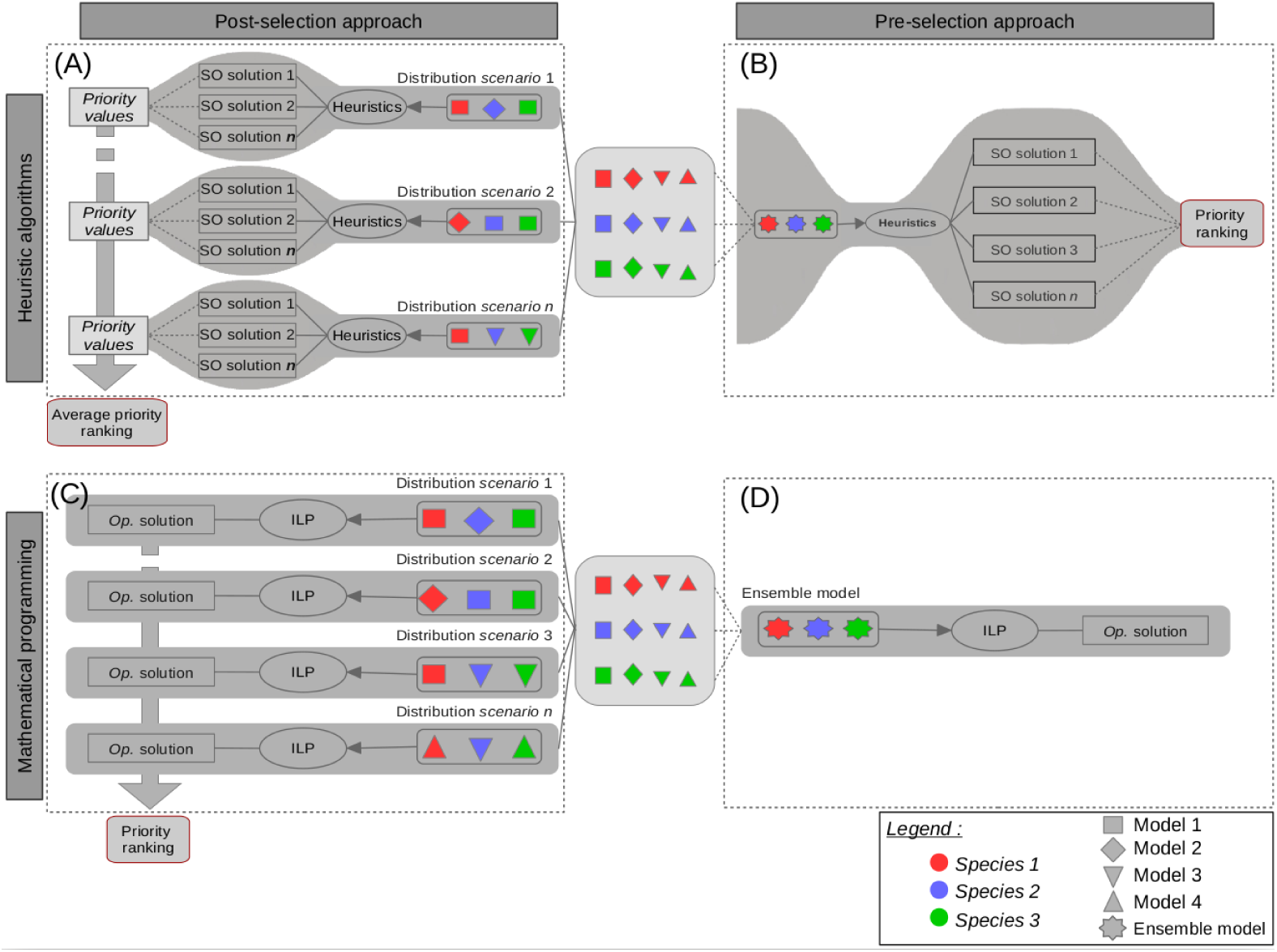
Schematic representation of four different approaches for the definition of spatial priorities when accounting for distribution modeling uncertainties. For each species (represented by different colors), distribution data were predicted using various correlative methods (each represented by different shape), which associate the species presence data within surveyed areas with predictor variables. (A) Following the post-selection approach, a set of sub-optimal solutions are generated for each distribution scenario, constructed by selecting randomly a model among all SDMs for each species. Average priority ranking values are calculated based on the priority ranking corresponding to each “distribution scenario”. (B) Following the pre-selection approach, distribution models are summarized into an ensemble model *a priori* to reserve selection. Priority ranking values are estimated for a set of sub-optimal solutions provided using a heuristic algorithm. (C) Following the post-selection approach, combined with the use of an exact optimization algorithm, priority ranking values are estimated for a set of optimal solutions, each corresponding to a different distribution scenario. (D) Following the pre-selection approach, combined with the use of a heuristic optimization algorithm, a unique optimal solution is generated based on ensemble models. The spread of uncertainties related to both models accuracy and solution efficiency is represented by grey shade.

Though the approach has never been implemented, we believe that mathematical programming optimization combined with a post-selection of SDM outputs leads to a reserve-selection framework that might provide flexibility while simultaneously addressing solution efficiency, thus avoiding the tacit trade-off existing between the two concepts. To exemplify the approach we analyzed an experimental design crossing pre- and post-selection of SDM outputs versus heuristics and exact mathematical optimizations (Figure 1). We used the Mediterranean Sea as a biogeographical template for our analyses, integrating the outputs of 8 SDM techniques for 438 fishes species.

## II. METHODS

### Species distribution modeling

The occurrences of species (i.e. geographic locations) used for SDMs were obtained from multiple databases: OBIS (Ocean Biogeographic Information System), Global Biodiversity Information Facility (GBIF), iNaturalist (A Community for Naturalists), VertNet (vertebrate biodiversity networks), Ecoengine (UC Berkeley’s Natural History Data) and Fishmed (Albouy et al., 2015). We filtered these data by removing potentially erroneous occurrences (e.g. located on continents) and keeping only the observations made since 1975. For the calibration of SDMs, temperature and salinity climatologies were acquired from the global World Ocean Database 2013 Version 2 at a spatial resolution of 1/4 °. These climatologies represent decadal averages of temperatures and salinities for 1975-1984, 1985-1994, 1995-2004 and 2005-2012 distributed over 40 standard depth layers. These data were interpolated at a spatial resolution of 1/12th of a degree (5 arcmin) and aggregated vertically by calculating mean values in the first 50 and 200 meters depth, for the calibration of pelagic and benthopelagic species respectively; and the last 50 meters depth for benthic and demersal species.

A set of 8 statistical correlative methods based on presence/pseudo-absence were used. These statictical methods belong to four main categories: multiple regressions (Generalized Linear Model, Generalized Additive Model, Multiple Adaptive Regression Splines), regression trees (Boosted Regression Tree, Random Forest, Classification Tree Analysis), Flexible Discriminant Analysis and Artificial Neural Network. These algorithms have been implemented using the BIOMOD2 multi-model platform (Thuiller et al., 2016). All models were evaluated using a cross-validation procedure with random 3-fold partitioning of occurrences data. For each cross-validation subset, 75% of occurrences are used for calibration and the remaining 25% for validation. In total, we obtained 24 suitability maps for each species, which were transformed into presence/absence maps by using the probability threshold that maximizes the models’ True Skill Statistic (TSS; Allouche et al., 2006). This criterion evaluates the predictive power of the models, taking into account both omission and commission errors, and indicates perfect agreement when close to one and a performance no better than random when close to zero or less.

We quantified the uncertainty in model predictions at two levels: species and assemblages. First for each species and within each planning unit, we calculated the average of binary predictions (i.e. models committee averaging; Thuiller et al. 2016). A committee averaging score close to 0 or 1 means that all models agree to predict 0 and 1 respectively. A score of 0.5 means that half the models predict 1 and the other half 0, and reflects a maximal prediction uncertainty. We then simplified these scores to measure the uncertainty in model predictions such that 0 represents a total agreement among models and 1 represents the situation where the same number of models predict an absence and a presence, using the following formula:

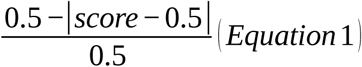

Second, at the assemblage level, we averaged the uncertainty scores for all species present in a grid cell to obtain an uncertainty map of SDM predictions.

### Species distribution scenarios and conservation targets

Only models with TSS greater than 0.6 were selected to be used for the establishment of various distribution datasets. First, SDM uncertainties are accounted for by summarizing the full range of predictions into an ensemble model, *a priori* to reserve selection, following the pre-selection approach. Here, for each species, several distribution maps were combined into one final ensemble by using averaging the predictions of selected SDMs. For the post-selection approach, we produced 100 different “distribution scenarios” by randomly sampling among the available distribution maps for each species.

We defined conservation targets, for each delivered dataset, such that restricted-range species have more ambitious objectives (Guilhaumon et al., 2015; Rodrigues et al., 2004; Venter et al., 2018). A target of 100% representation has been set for species with restricted distribution (range <1000 km2) and a target of 10% has been used for widespread species (those with a geographic extent exceeding the two-thirds of all species ranges). For species with intermediate-sized ranges the target was interpolated as a linear function of the log-transformed range (Rodrigues et al, 2004). In addition, we modified conservation targets based on the threat level of the species as determined by the IUCN Red List categories (Abdul Malak et al., 2011). Specifically, conservation targets for critically endangered species were promoted to protect 100% of their spatial range.

### Reserve selection algorithms and priority rankings

We identified conservation solutions that ensure species target achievement while minimizing the total cost of selected Planning Units (PUs or sites), following the the minimum set problem formulation (Moilanen and Ball, 2009). We used Integer Linear Programming as an implementation of exact optimization (ILP; Figure 1, C-D; Possingham et al., 2000); and the Marxan decision support tool to implement heuristic reserve-selection (Figure 1, A-B).

The ILP problem was formulated to minimize the cost of the solution (the objective function) while respecting a set of linear constraints:

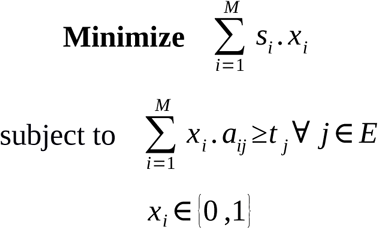

Where M is the number of PUs, and E is the set of species. Let *a*_*ij*_ =1 if the species (*j* ∈ *E*) is present in the PU (*i* ∈ *S*) and zero otherwise. The boolean variable *x*_*i*_=1 if the site (*i* ∈ *S*) is selected and zero otherwise. Each PU (*i* ∈ *S*) is described by it’s surface area *s*_*i*_. *t* _*j*_ is the minimum amount of each species range to be included in the solution (i.e. the conservation target of species (*j* ∈ *E*)). The ILP problems were solved using the GUROBI software, which implements the “branch and bound” exact algorithm (Gurobi Optimization, 2012).

Marxan (Ball et al., 2009) uses a simulated annealing algorithm (a meta-heuristic algorithm) to identity sub-optimal systems of priority areas. This is done by minimizing iteratively an objective function summing the total cost of PUs in the solution and penalties for not met species targets (Species’ Penalty Factor, SPF, here set such as to ensure the representation of all species). We set the boundary length modifier to 0 (a parameter that measure the trade-off between cost and compactness of the solution), as our aim was to examine differences in the selection of priority areas among the scenarios and not to design an MPA network with a desirable level of compactness. In our study, Marxan was run 100 times and consisted of 1,000,000 iterations per run. In this application, the cost of each site was equal to its area, favoring the selection of sites with high ecological importance.

A priority ranking for planning units was calculated as the selection frequency of each site across a number of conservation solutions (optimal or sub-optimal), so then sites selected above a certain threshold-percentage of solutions are considered as high-priority conservation areas (selected more than 90 times out of the 100 solutions; Andelman and Willig, 2002; Leslie et al., 2003). We first calculated the PUs selection frequency across the 100 sub-optimal solutions derived from Marxan outputs following the pre-selection approach (Figure 1, B). Second, following the post-selection approach, we calculated selection frequency over the 100 optimal solutions, each based on a different “distribution scenario” (Figure 1, C). Finally, the set of 100 sub-optimal solutions was used to calculate selection frequencies, each based on a different distribution scenario. Those values were summarized by averaging selection frequencies, and provide a final average priority ranking as outcome of the post-selection approach with heuristic reserve selection (Figure 1, A).

### Comparative analysis

First, we compared the distributions of priority rankings, derived from the pre-selection (coupled with heuristic reserve selection algorithm) and post-selection (coupled with exact reserve selection algorithm) approaches, using 10-bins histograms. As histograms give no insight into the spatial location of the differences and similarities between priority ranking values, we mapped the differences in PUs ranking and measured the overlap in high-priority conservation areas between the two approaches. Besides, the Wilcoxon test (t-test for paired samples), was used to determine statistically significant differences between the PU rankings obtained. These rankings were compared to the uncertainties associated with the modeling of species distributions using Spearman’s correlation test, to determine whether these uncertainties are better represented by one approach or another.

Additionally, we compared the total protected area for each conservation solution obtained, based on the different approaches, and quantified the efficiency of heuristics outcomes for each “distribution scenario” (i.e. gap to optimality) as the difference in total area with respect to the optimal solution.

## II. RESULTS

Overall, SDMs showed variable performances in predicting observed species distributions, with TSS values ranging from 0 to 0.98 (0.73 *±* 0.1; *mean± standard deviation*). To avoid spurious conclusions based on unfair predictions, only models with TSS greater than 0.6 were used for the remaining analyses. For all 438 species considered, several statistical methods showed good performance in predicting observed distributions, which prevented a single “best” model from being distinguished.

In addition, significant variability was observed in the modeled distribution ranges of species (Figure A2.1). The inferences about species distributions and, consequently, conservation targets were decisively dependent on the SDMs modeling approach, with substantial uncertainties associated with the choice of a single best statistical modeling technique. This species-level variability has resulted in spatially-structured uncertainties at the assemblage level (Eq.1), with congruence areas mainly located in pelagic environments; and areas of disagreement at the edge of the species distribution areas, along the margins of the continental shelf, notably along the shores of the Aegean and Ionian seas (Figure 2).

**Figure 2:**
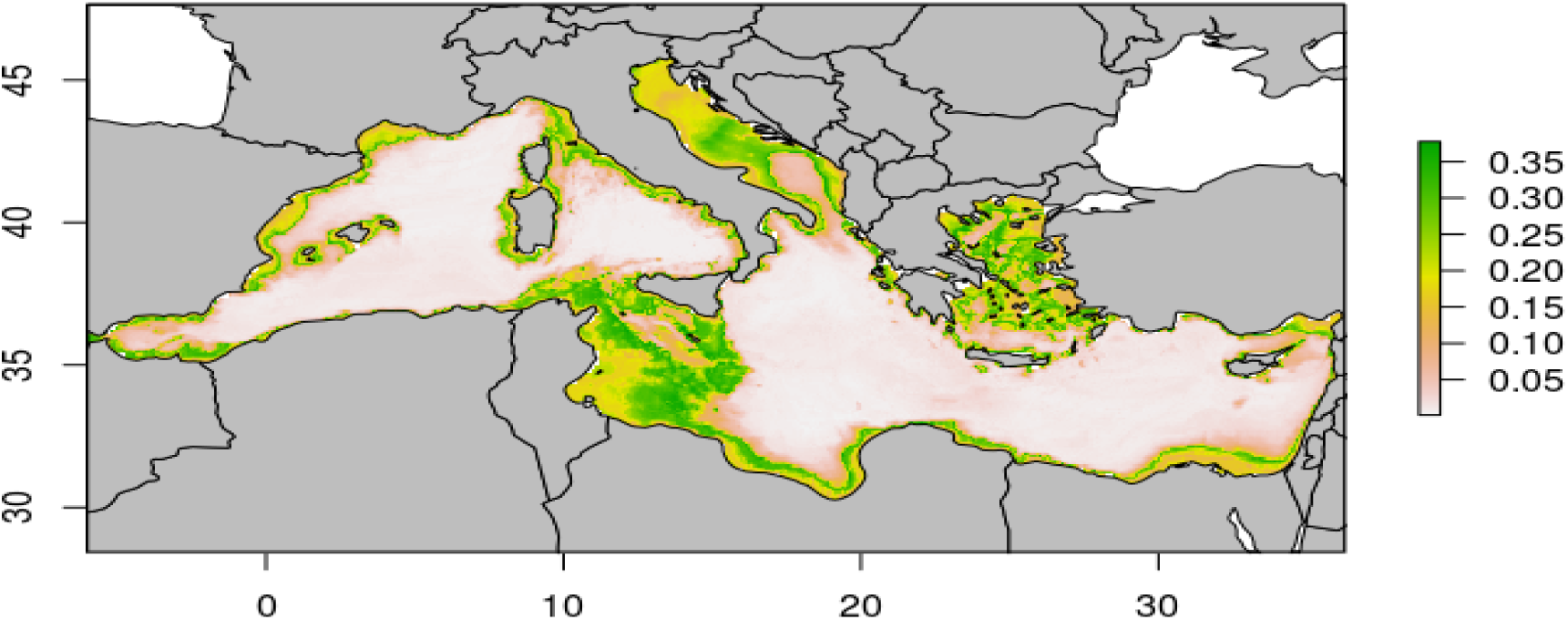
Map of uncertainties in SDM predictions. Uncertainty calculated at the species level (eq. 1) and averaged across species present in each grid cell. Green areas indicate cells for which model predictions uncertainties are the highest.

The uncertainties of the SDMs have often been taken into account, in the literature, by opting for the pre-selection approach, based on a consensual overall model as input data for SCP exercises (Appendix 1). The conservation results based on this pre-selection approach, with the ensemble model as input data for Marxan, revealed that the sub-optimal solutions contained 2.85 to 3.34% more PUs than the optimal solution, generated using ILP. We identified 32.6% of irreplaceable PU’s that were consistently selected across the 100 sub-optimal solutions (Fig. A2.2-b), while the minimum area required to achieve species targets reached 36.7% of PUs identified as the optimal solution (Fig. A2.2-a). Indeed, The distribution of PU’s selection frequency was strongly bimodal using both exact and heuristics algorithms, with most of the PUs belonging to the first and last 20% quantiles (0-20 and 80-100; Figure 3).

**Figure 3:**
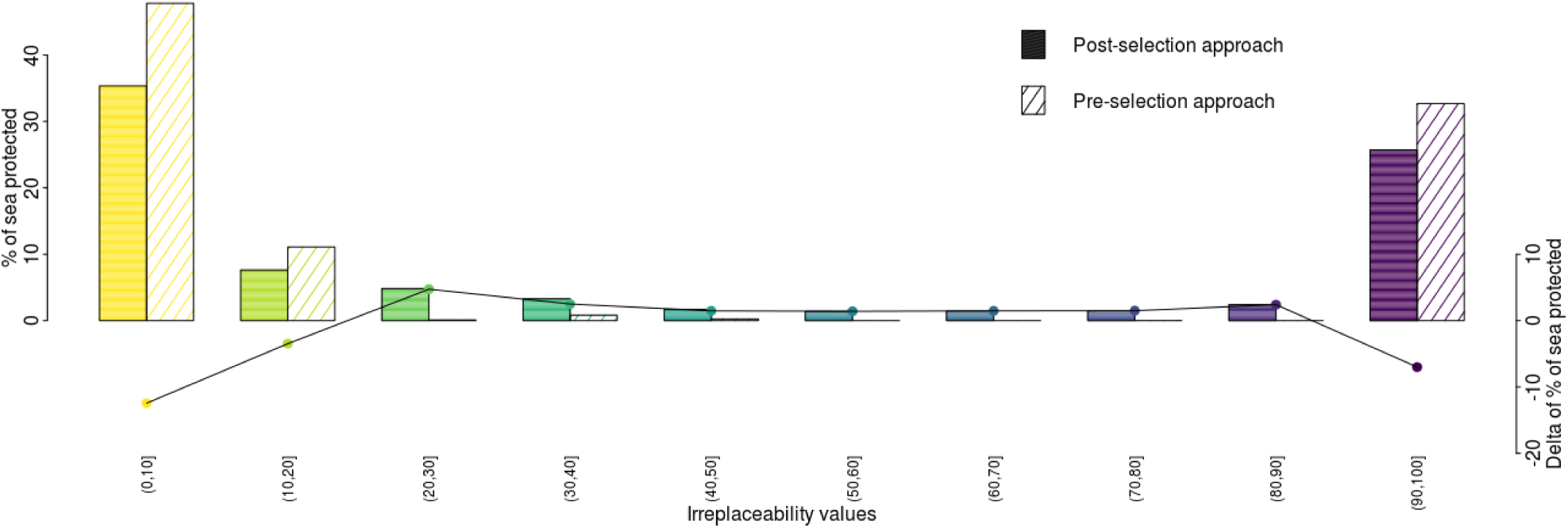
Frequency distributions of priority ranking values derived from two approaches used for integrating species distribution uncertainties. Bars of full color represents the result of post-selection approach while hatched bars represent result derived following the pre-selection approach. The curve represents the frequency differences across different levels of priority between two approaches.

Following the post-selection approach, 100 “distribution scenarios” were generated by randomly sampling one model for each species among the SDMs selected to have a good predictive performance, thus considering the full range of good predictions (Figure 1, A-C). Then, we identified conservation solutions and investigated the variability in the network size and in the spatial distribution of the selected PUs. Indeed, the optimal solutions, each based on a different distribution scenario, vary in terms of the area of the conservation solutions, ranging from c. 27% to c. 43% of the Mediterranean Sea (Figure A3.2-a). Among those optimal solutions, 85% of PUs were selected at least once. Selection frequencies derived from the post-selection approach display increasingly more fully flexible PUs, allocating moderate values (20 to 80) to a greater number of Pus than the pre-selection approach (Figure 3). With this respect, the number of high priority PUs were about 7% lower than for the pre-selection approach (Figure 3). Moreover, the Wilcoxon test, used to compare the priority rankings of the UPs derived from the two pre-selection and post-selection approaches, was significant (p-value<0.5), revealing that the selection frequency of the PUs depends on the strategy followed for considering species distribution uncertainties.

Further, we examined the spatial mismatch of selection frequencies between two approaches (Figure 4). Only 14 % of areas were identified as totally irreplaceable by both approaches (i.e. selection frequency of 100 %; Figure 4). In contrast, the frequency selection of 9.83% of planning units showed differences of 20% and more between the two approaches. Areas with higher priority following the post-selection approach, are mainly located along the Aegean Sea and Ionian Sea coasts; while areas presenting higher priority following the pre-selection approach are spread in small and isolated areas along all Mediterranean coasts (Figure 4). The correlation between the SDMs uncertainties and the selection frequency values derived from the post-selection approach was greater than that of the pre-selection approach (Spearman r_s_ 0.77 and 0.57 respectively). Hence, the conservation solutions identified with the post-selection approach were more representative of the uncertainty map of predictions than the pre-selection solutions.

**Figure 4:**
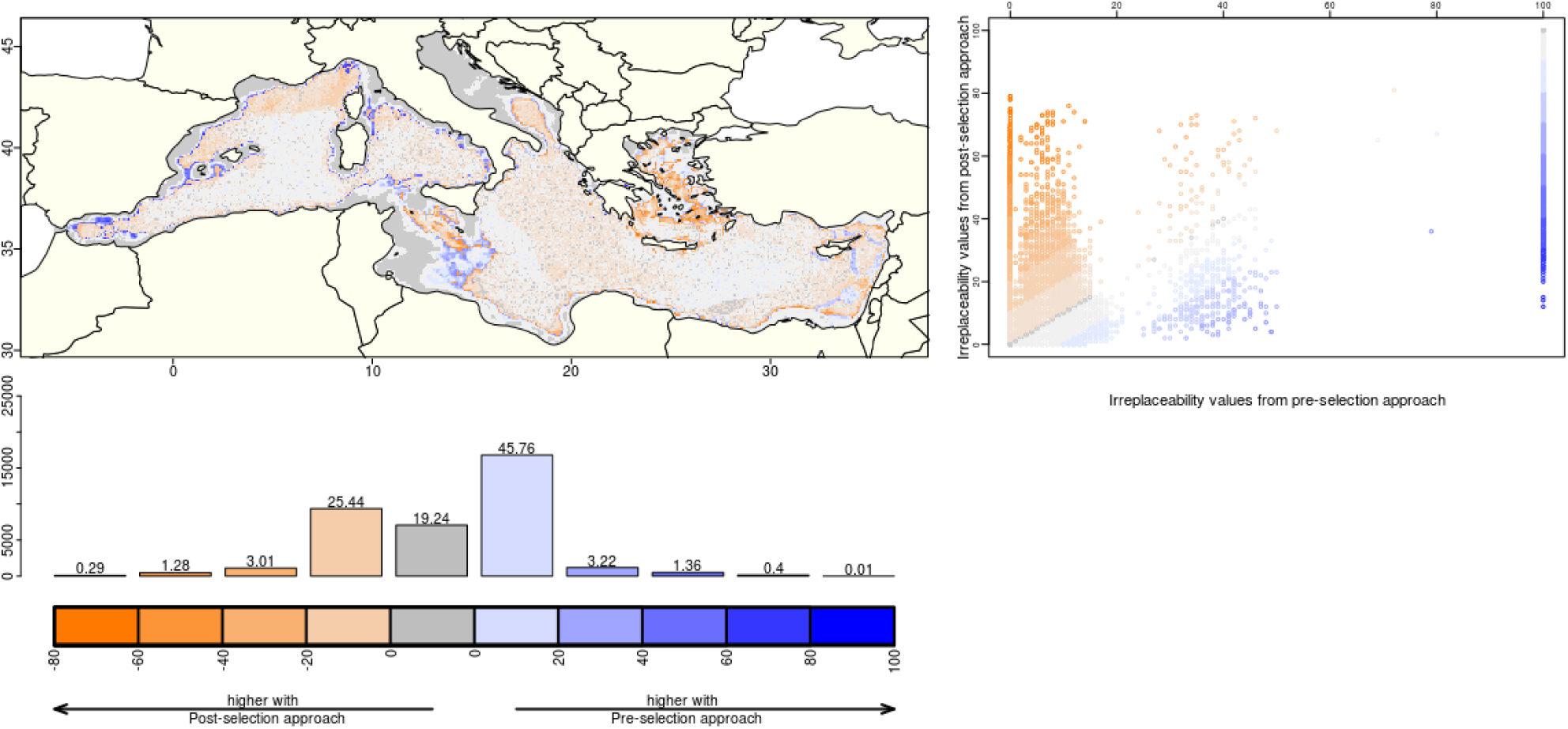
Comparison of priority ranking values based on two approaches including uncertainty associated with distribution modeling. (a) map of difference in PUs selection frequency (b) scatter plot showing the PUs selection frequency under the different approaches. (c) Histogram of selection frequence differences.

The post-selection approach can also be based on sub-optimal solutions, derived from heuristics reserve selection algorithms (Figure 1, A). In this study, the statistics on the optimality gaps, calculated between the optimal and sub-optimal solutions for the each “distribution scenario”, reveal that half of the solutions identified using heuristics are c. 3 to 8% more costly than those found by the exact optimization (Figure 5).

**Figure 5:**
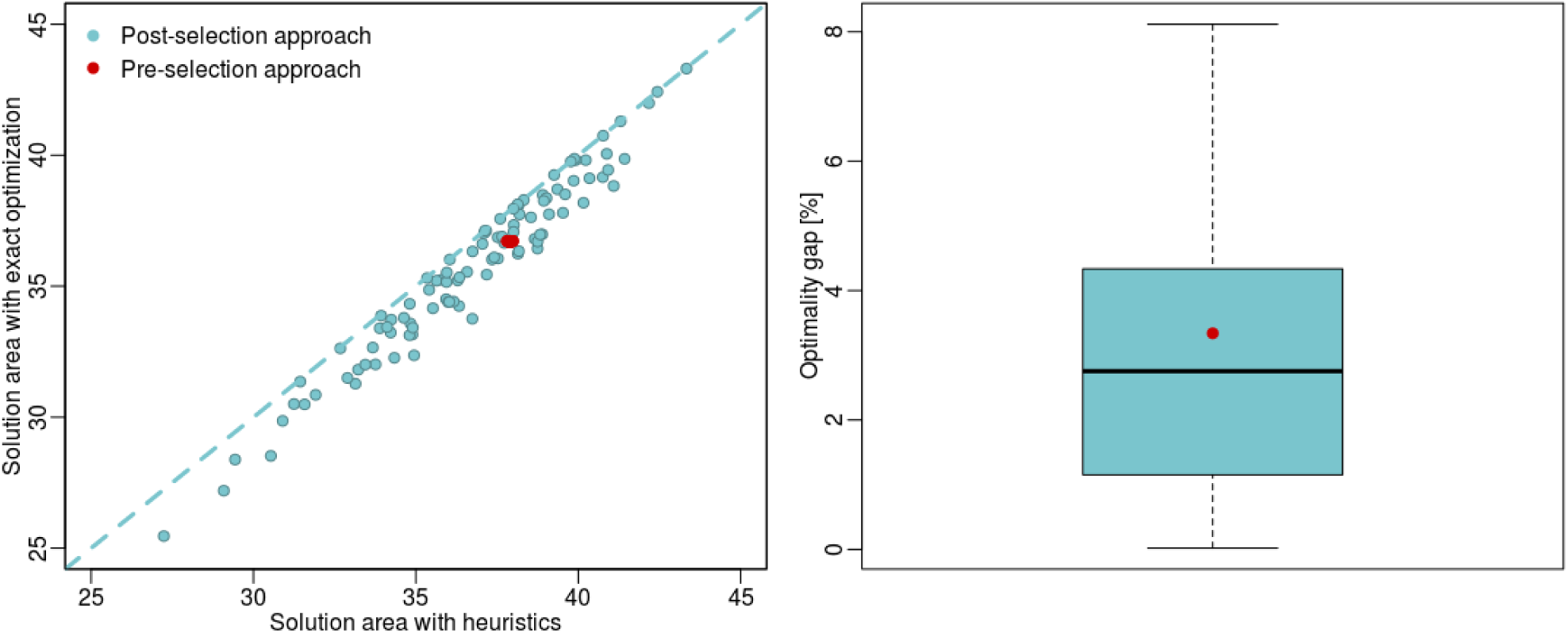
(a) Comparison of protected areas expressed as percentages of the sea, between heuristics and optimal conservation outcomes for both pre-selection and post-selection approaches; Each point represents the % of protected areas necessary to achieve species target. (b) Boxplot of optimality gap between sub-optimal and optimal conservation outcomes for each “distribution scenario”. The gap between best-solution and optimal solution based on pre-selection approach is shown by the red point.

## III. Discussion

Systematic conservation planning is a data-intensive process, requiring species occurrences data that cover the overall landscape (Rondinini et al., 2006; Tulloch et al., 2016). Such knowledge about species distributions is usually so scarce and geographically variable that conservation applications rely increasingly on modeling techniques to predict the distribution of species in unsurveyed areas (Appendix 1). The literature review revealed that the choice of the statistical method used to model distributions is likely to be opportunistic in conservation applications as the great majority of studies identified conservation solutions on the basis of species distributions modelled with only one method (Appendix A1). Using a single and a priori chosen model, particularly for rare species, may yield spurious predictions, affecting the selection of MPAs and jeopardizing the adequacy of conservation targets and the representativeness of selected MPAs (Guillera-Arroita et al., 2015; Pearson et al., 2006). Before any conservation exercise, the predictive performance of SDMs should thus be evaluated, using evaluation metrics (here TSS). In this study, various models yielded equivalent predictions performances, which prevented to distinguish a single “best” model. It is thus crucial for any conservation exercise to start by testing different statistical methods for species distribution modeling, and to evaluate the predictive performance of the models obtained (Molloy et al., 2016).

The uncertainty related to the choice of an SDM resulted in substancial variability in species predicted geographic ranges and conservation targets. It was spatially translated into spatially structured uncertainties by identifying areas where different SDMs agree/or disagree in predicting species presence or absence. The spatial difference between predictions depends on the accuracy of the SDMs, and the marginality and specialization of the niches of the species (Zhang et al., 2015). The selection of incongruent PUs is risky for the representativeness of the conservation solution, because the species presence may not be certain, according to the statistical method used for modeling their niches. Spatial uncertainties, therefore, need to be given more attention before making any decisions based on predictions, notably for species with high marginality and niche specialization.

Those modeling uncertainties pertain in the literature, and most studies opt for the pre-selection approach in order to address them (Appendix 1; e.g. Alagador et al., 2016; Bush et al., 2014; Faleiro and Loyola, 2013). The use of an exact reserve selection algorithm to solve an ILP problem based on predictions from the ensemble models, reveals a unique, optimal and efficient conservation solution, covering 36.7% of the Mediterranean Sea. On the contrary, Marxan’s heuristic algorithm produces a set of sub-optimal solutions, thus making it possible to estimate the priority ranking of PUs. However, the ranking obtained is bimodal, with 32.6% of the Mediterranean Sea completely irreplaceable to achieve the conservation objectives. Actually, this potential for replacement of PUs, as members of protected area systems, increased with an increasing total area of sites needed to achieve the representation targets (Pressey et al., 1996). This means that flexibility obtained following the pre-selection approach depends upon the inherent variability of optimality gaps, and therefore on the algorithm’s ability to generate effective solutions (here 2.85 to 3.34 % more sites than the optimal solution).

Flexibility could also be provided by applying the algorithms repeatedly under different conditions (Rebelo and Siegfried, 1992). Following the post-selection approach, different “distributions scenarios” are considered by randomly sampling model predictions for each species among the full range of predictions (Figure 1, A-C). The variability in species predicted geographic ranges resulted in differences in term of the covered areas proportional to the mediterranean sea across optimal solutions, each based on a different distribution scenario. This approach produces values of priority ranking significantly different from the values generated following the pre-selection approach, supporting the fact that conservation decisions based on predicted distributions are sensitive to the approach used to summarize the modeling uncertainties (Meller et al., 2014). The post-selection approach generated priority ranking values with more replaceable UPs and greater flexibility to decision-makers than the pre-selection approach. Those ranking values are shown to be more representative of distribution uncertainties than the pre-selection outcomes. Hence, areas with higher priority following the post-selection approach (mainly located along the Aegean sea and Ionian sea coasts, and the continental shelf margins of the eastern Mediterranean sea), highlights areas with low SDMs precision or/and comprising species with high marginality and niche specialization. The post-selection approach represents thus fundamentally the most accurate way of site prioritization, for highlighting areas with spatial discrepancies between different SDMs.

Representing the data uncertainties, as a base for the negotiation of conservation planning, is of greater concerns in the literature when planning for climate change adaptation (Appendix 1). Apart from producing a ranking of priorities related to the capabilities of the algorithm to find the most effective solution, the pre-selection approach is based on an ensemble model that does not necessarily provide a more accurate future projection than another niche model potentially realizable by the species (Zhang et al., 2015).

In practice, this approach can be based on optimal solutions, but also the best sub-optimal solutions from heuristics. Several studies have commented on the sub-optimality of conservation solutions (e.g. Moilanen, 2008; Onal and Briers, 2002; Pressey et al., 1996). As matter of fact, Vanderkam et al. (2007) found that heuristics yielded protected area systems that were 2%–70% larger than the network identified by an optimal algorithm (i.e. Mathematical programming), depending on the formulation of optimization problem and the heuristic algorithm used. It should also be noted that heuristic algorithms could not inform the user of the degree of solutions sub-optimality. Our results show that the sub-optimality between marxan’s and optimal solutions for the same “distribution scenario” can reach up to 8% of the study area. Moilanen (2008) concludes that a 5%–10% efficiency loss out of limited resources can be considered meaningful. Combine with the fact that over the 10% of marine areas targeted for protection by CBD only 7.14% of the Mediterranean sea is under a legal conservation designation (MAPAMED, 2018), these results reinforce the need to opt for an algorithm that saves scarce resources allocated to conservation.

To conclude, our study highlighted that the modeling technique is an important factor to consider when using SDMs as inputs for SCP exercises. The priority ranking values estimated following two strategies for accounting distribution modeling uncertainties can be significantly different, leading to distinct perspectives on prioritization of conservation actions, and jeopardizing the adequacy and representativeness of conservation outcomes. While the post-selection approach estimates sites priorities by varying solutions efficiency, the post-selection approach (coupled with the exact algorithm) provides decision-makers with the greatest information related to the uncertainties in species distributions knowledge across different modeling techniques. This information is communicated as priority values, where the most irreplaceable areas are selected across different potential “distribution scenarios”. Thus, the uncertainty associated with predictions needs to be documented and to be effectively communicated to managers, e.g. by following post-selection approachs and the selection of MPAs must be optimal and efficient, e.g. based on an exact optimization algorithm. This approach provides a foundation for a conservation planning that respects the fundamental principles of SCP, aiming for adaptive and effective management of conservation resources.

