## Appendix for "Are trade-offs between flexibility and efficiency in systematic conservation planning avoidable ?"

### <u>Appendix 1</u>

In order to assay main trends on the use of SDMs through spatial conservation planning, a literature search was conducted using Web of Science (WoS). The intent of the review is to assess the ways to which SDMs have been integrated into conservation planning by evaluating two questions: 1) What types of models have been included ? and 3) How uncertainty between different models has been accounted for in conservation planning ?

To identify relevant literature, we carried out a topic search examining the title, keywords and abstracts among papers, using the folowing query :

TS = ( ("species distribution model*" OR "habitat model*" OR "ecological niche model*" OR "habitat suitability ind*" OR "habitat suitability mod*" OR "occupancy mod*" OR maxent OR "presence-absence mod*" OR "presence-only mod*" OR "niche model*" OR "climate change" OR "changing climate") <u>AND</u> ("Spatial planning" OR "Spatial optimization" OR "Marine reserve*" OR "spatial prioritization" OR "reserve selection" OR "area-selection algorithm*" OR "spatial conservation prioritization" OR "conservation plan*" OR "land use plan*" OR "regional plan*") <u>AND</u> ( ResNet OR worldmap OR zonation OR "C-plan" OR "Marxan" OR "linear programming" OR "mathematical optimization") )

This query results initially in 137 papers. Articles without study case for both distribution modeling and conservation planning application were excluded. Our literature review exclude further national journals, reviews, case reports, letters, editorials, and conference abstracts. Furthermore, we browsed the most frequently cited papers in the gathered corpus , to have a final corpus composed of 76 articles in total. Still, our review may be incomplete, it remains representative.

We then extracted the following informations from all studies, provided in table A1 (when details were available) : (1) Year of publication (2) Number of techniques used for species distribution modeling (3) projection in futur under climate change scenarios (Boolean) (4) reserve-selection algorithm used for conservation planning (scoring ; heuristics or mathematical programming) (5) Uncertainty of feature data (deal with it or not ; Boolean).

The use of SDMs is increasingly favored and more common in SCP applications (Figure A1). Most conservation applications have neglected the uncertainties associated with different modeling techniques, by using a unique modeling method, chosen *a priori*, to predict the distribution of species (n = 65); while others have examined a wide range of techniques and have used only the “best” model identified in terms of predictive performance (Table A1; e.g. Leach et al., 2013;

Passoni et al., 2017; Walther and Pirsig, 2017). However, when such particular solutions are evaluated using other statistical method, the representation of species in the conservation solution might not meet the required target level (Loiselle et al., 2003).

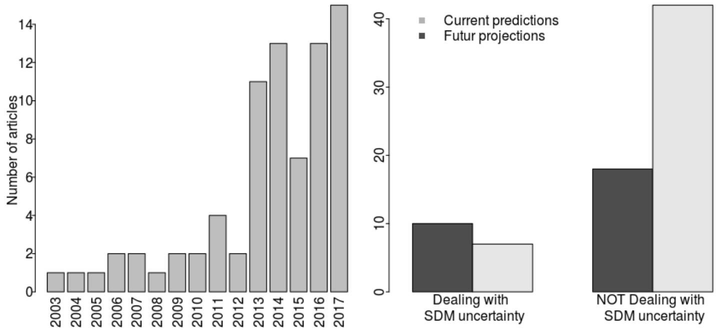


**Figure A1 :** Results of the analysis of 76 publications generating conservation priorities, based on the distribution patterns of species: the left, the number of publications per year; On the right, the number of publications according to whether or not they take into account the uncertainties related to modeling methods, for current and future predictions.

Only c. 23% of the reviewed publications incorporated SDMs uncertainties explicitly into the conservation planning process, especially for the modeling of future projections (Figure A1). To do this, all studies have adopted ensemble prediction approaches (called ensemble or multi-model models or model averaging), representing a consensus model with median or mean prediction values, in order to combine different SDMs, and significantly improve the precision of the predicted distributions (e.g. Alagador et al., 2016, Faleiro and Loyola, 2013). This approach, hereafter referred to as the pre-selection approach, aims to reduce uncertainty a priori to conservation planning, so that the distributions predicted by the ensemble models are used as inputs for the optimization algorithms. Meller et al. (2014) introduced another approach, hereafter referred to as the post-selection approach, in which different "distribution scenarios" are constructed by randomly sampling one model from across the range of SDMs; and used heuristic algortithm to identify conservation priorities.

The effectiveness of conservation solutions has been less taken into account when SDMs are involved in the SCP process. Most conservation applications (about 80% of the reviewed publications) used heuristic algorithms, with slightly more than half used Zonation (n = 42; Moilanen et al., 2009) and almost one quarter used Marxan (n= 19; Ball and Possingham, 2000). Only 4% (n = 3) of the reviewed publications implemented mathematical programming to provide optimal conservation solutions. All three studies have a pre-selection approach for managing SDMs uncertainties.

**Table A1 : List of publications identified in the review**

| N° | Title | Authors and year of publication | Journal of publication | Number of distribution modeling techniques | Model used for SCP (- / Ensemble / Best) | Future projections | Reserve selection algorithms | Dealing with SDMs uncertainties |
|---|---|---|---|---|---|---|---|---|
| 1 | Planning for conservation and restoration under climate and land use change in the Brazilian Atlantic Forest | (Zwiener et al., 2017) | DDI | 1 | - | Yes | Zonation | Yes |
| 2 | The role of macroinvertebrates for conservation of freshwater systems | (Nieto et al., 2017) | EE | 1 | - | No | Zonation | No |
| 3 | Trade-offs in carbon storage and biodiversity conservation under climate change reveal risk to endemic species | (Reside et al., 2017) | Biological Conservation | 1 | - | Yes | Zonation | Yes |
| 4 | Integrating biodiversity, ecosystem services and socio-economic data to identify priority areas and landowners for conservation actions at the national scale | (Di Minin et al., 2017) | Biological Conservation | 1 | - | No | Zonation | No |
| 5 | Analysis of Trade-Offs Between Biodiversity, Carbon Farming and Agricultural Development in Northern Australia Reveals the Benefits of Strategic Planning | (Morán-Ordóñez et al., 2017) | Conservation letters | 1 | - | No | Zonation | No |

| | | | | | | | | |
|---|---|---|---|---|---|---|---|---|
| 6 | Prioritizing Management of the Invasive Grass Common Reed (Phragmites australis) in Great Salt Lake Wetlands | (Long et al., 2017) | Invasive Plant Science and Management | 1 | Ensemble | No | MCA | No |
| 7 | Assessing the effectiveness of China's protected areas to conserve current and future amphibian diversity | (Chen et al., 2017) | DDI | 3 | Ensemble | Yes | Notation | No |
| 8 | Dealing with cumulative biodiversity impacts in strategic environmental assessment: A new frontier for conservation planning | (Whitehead et al., 2017) | Conservation letters | 1 | - | No | Zonation | No |
| 9 | Determining conservation priority areas for Palearctic passerine migrant birds in sub-Saharan Africa | (Walther and Pirsig, 2017) | Avian | 9 | Best | No | Marxan | No |
| 10 | Spatial conservation prioritization for dominant tree species of Chinese forest communities under climate change | (Wan et al., 2017) | Climatic Change | 1 | Ensemble | Yes | Zonation | No |
| 11 | From pest data to abundance-based risk maps combining eco-physiological knowledge, weather, and habitat variability | (Lacasella et al., 2017) | Ecological Applications | 1 | - | No | Zonation | No |
| 12 | Framework for strategic wind farm site prioritisation based on modelled wolf reproduction habitat in Croatia | (Passoni et al., 2017) | Eur J Wildl res | 1 | Best | No | Marxan | No |

| | | | | | | | | |
|---|---|---|---|---|---|---|---|---|
| 13 | Predicted distribution of whales at risk: identifying priority areas to enhance cetacean monitoring in the Northwest Atlantic Ocean | (Gomez et al., 2017) | Endangered Species Research | 1 | - | No | Notation | No |
| 14 | Incorporating dynamic distributions into spatial prioritization | (Runge et al., 2016) | Ddi | 1 | - | No | Marxan | No |
| 15 | Species distributions represent intraspecific geneticdiversity of freshwater fish in conservation assessments | (Hermoso et al., 2016) | Freshwater Biol. | 1 | - | No | Marxan | No |
| 16 | Habitat mapping as a tool for water birds conservation planning in an arid zone wetland: The case study Hamun wetland | (Maleki et al., 2016) | Ecological Engineering | 1 | - | No | Notation | No |
| 17 | Synergistic effects of climate and land-use change on representation of African bats in priority conservation areas | (Smith et al., 2016) | Ecological Indicators | 1 | - | Yes | Zonation | Yes |
| 18 | Integrating Climate Change Resilience Features into the Incremental Refinement of an Existing Marine Park | (Davies et al., 2016) | Plos One | 1 | - | No | Marxan | No |
| 19 | Identifying areas of optimal multispecies conservation value by accounting for incompatibilities between species | (Beaudry et al., 2016) | Ecological Modelling | 1 | - | No | ILP | No |

| | | | | | | | | |
|---|---|---|---|---|---|---|---|---|
| 20 | Climate change, species range shifts and dispersal corridors: an evaluation of spatial conservation models | (Alagador et al., 2016) | MEE | 9 | Ensemble | Yes | ILP | No |
| 21 | Large expansion of oil industry in the Ecuadorian Amazon:biodiversity vulnerability and conservation alternatives | (Lessmann et al., 2016) | EE | 1 | - | No | Marxan | No |
| 22 | Downscaling patterns of complementarity to a finerresolution and its implications for conservation prioritization | (de Albuquerque and Beier, 2016) | EE | 1 | - | No | Zonation | No |
| 23 | Optimising different types of biodiversity coverage of protected areas with a case study using Himalayan Galliformes | (Dunn et al., 2016) | Conservation Biology | 1 | Ensemble | No | Zonation | Yes |
| 24 | Reserve network planning for fishes in the middle and lower Yangtze River basin by systematic conservation approaches | (Huang et al., 2016) | Science China | 1 | - | No | Marxan | No |
| 25 | Incorporating natural and human factors in habitat modelling and spatial prioritisation for the Lynx lynx martinoi | (Laze and Gordon, 2016) | Web ecol | 1 | - | No | Zonation | No |
| 26 | Identifying appropriate protected areas for endangered fern species under climate change | (Wang et al., 2016) | SpringerPlus | 1 | - | Yes | Zonation | No |

| | | | | | | | | |
|---|---|---|---|---|---|---|---|---|
| 27 | Phylogenetic diversity meets conservation policy: small areas are key to preserving eucalypt lineages. | (Pollock et al., 2015) | Phil. Trans. R. Soc. | 1 | - | No | Zonation | No |
| 28 | Systematic site selection for multispecies monitoring networks | (Carvalho et al., 2016) | JAE | 1 | - | Yes | Marxan | No |
| 29 | Developing conservation strategies for Pinus koraiensis and Eleutherococcus senticosus by using model-based geographic distributions | (Wan et al., 2016) | Journal of forestry | 1 | - | Yes | Marxan | Yes |
| 30 | Spatial conservation priorities are highly sensitive to choice of biodiversity surrogates and species distribution model type | (Lentini and Wintle, 2015) | Ecography | 1 | - | No | Zonation | Yes |
| 31 | Conservation of future boreal forest bird communities considering lags in vegetation response to climate change: a modified refugia approach | (Stralberg et al., 2015) | DDI | 1 | - | Yes | Zonation | No |
| 32 | Supporting underrepresented forests in Mesoamerica | (de Albuquerque et al., 2015) | Natureza&Conse rvaceo | 1 | - | No | Zonation | No |
| 33 | Prioritizing Regions to Conserve a Specialist Folivore: Considering Probability of Occurrence, Food Resources, and Climate Change | (Adams-Hosking et al., 2015) | Conservation letters | 1 | - | Yes | Zonation | No |

| | | | | | | | | |
|---|---|---|---|---|---|---|---|---|
| 34 | Combined Use of Systematic Conservation Planning, Species Distribution Modelling, and Connectivity Analysis Reveals Severe Conservation Gaps in a Megadiverse Country (Peru) | (Fajardo et al., 2014) | Plos One | 1 | - | No | Marxan | No |
| 35 | Freshwater conservation planning under climate change: demonstrating proactive approaches for Australian Odonata | (Bush et al., 2014) | JAE | 9 | Ensemble | Yes | Marxan | No |
| 36 | Integrating Biological and Social Values When Prioritizing Places for Biodiversity Conservation | (Whitehead et al., 2014) | Conservation Biology | 1 | - | No | Zonation | No |
| 37 | Incorporating climate change in conservation planning for freshwater fishes | (Bond et al., 2014) | DDI | 1 | - | Yes | Zonation | No |
| 38 | Projecting current and future location, quality, and connectivity of habitat for breeding birds in the Great Basin | (Fleishman et al., 2014) | ECOSPHERE | 1 | - | Yes | Zonation | No |
| 39 | Model-based conservation planning of the genetic diversity of Phellodendron amurense Rupr due to climate change | (Wan et al., 2014b) | EE | 1 | - | Yes | Zonation | No |

| | | | | | | | | |
|---|---|---|---|---|---|---|---|---|
| 40 | Maximizing species conservation in continental Ecuador: a case of systematic conservation planning for biodiverse regions | (Lessmann et al., 2014) | EE | 1 | - | No | Marxan | No |
| 41 | Planning the priority protected areas of endangered orchid species in northeastern China | (Wan et al., 2014a) | BIODIVERSITY AND CONSERVATIO N | 1 | - | No | Zonation | No |
| 42 | Designing Optimized Multi-Species Monitoring Networks to Detect Range Shifts Driven by Climate Change: A Case Study with Bats in the North of Portugal | (Amorim et al., 2014) | Plos One | 1 | Ensemble | Yes | Marxan | No |
| 43 | Identifying minimal sets of survey techniques for multi-species monitoring across landscapes: an approach utilising species distribution models | (Bino et al., 2014) | INTERNATION AL JOURNAL OF GEOGRAPHIC AL INFORMATION SCIENCE | 1 | Best | No | Marxan | No |
| 44 | A spatial conservation prioritization approach for protecting marine birds given proposed offshore wind energy development | (Winiarski et al., 2014) | Biological Conservation | 1 | - | No | Zonation | Yes |
| 45 | Endemic wild potato (Solanum spp.) biodiversity status in Bolivia: Reasons for conservation concerns | (Cadima et al., 2014) | JOURNAL FOR NATURE CONSERVATIO N | 1 | - | No | SIG | No |

| | | | | | | | | |
|---|---|---|---|---|---|---|---|---|
| 46 | Shifting protected areas: scheduling spatial priorities under climate change | (Alagador et al., 2014) | JAE | 7 | Ensemble | Yes | PLNE | Yes |
| 47 | Socioeconomic and political trade-offs in biodiversity conservation: a case study of the Cerrado Biodiversity Hotspot, Brazil | (Faleiro and Loyola, 2013) | DDI | 9 | Ensemble | No | Zonation | Yes |
| 48 | Improving bioregional frameworks for conservation by including mammal distributions | (Bino et al., 2013) | AUSTRAL ECOLOGY | 1 | Best | No | Marxan | No |
| 49 | Modeling climate change impacts on tidal marsh birds: Restoration and conservation planning in the face of uncertainty | (Veloz et al., 2013) | ECOSPHERE | 1 | Best | Yes | Zonation | Yes |
| 50 | Egypt's Protected Area network under future climate change | (Leach et al., 2013) | Biological Conservation | 1 | Best | Yes | Zonation | Yes |
| 51 | The ice age ecologist: testing methods for reserve prioritization during the last global warming | (Williams et al., 2013) | GEB | 1 | - | Yes | Zonation | No |
| 52 | Using Seabird Habitat Modeling to Inform Marine Spatial Planning in Central California's National Marine Sanctuaries | (McGowan et al., 2013) | Plos One | 4 | Ensemble | Yes | Marxan | No |

| | | | | | | | | |
|---|---|---|---|---|---|---|---|---|
| 53 | A new spin on a compositionalist predictive modelling framework for conservation planning: A tropical case study in Ecuador | (Mateo et al., 2013) | Biological Conservation | 1 | - | No | Zonation | No |
| 54 | A preliminary assessment of the effectiveness of the Mesoamerican Biological Corridor for protecting potential Baird's tapir (Tapirus bairdii) habitat in southern Mexico | (Mendoza et al., 2013) | INTEGRATIVE ZOOLOGY | 3 | Ensemble | Yes | Zonation | No |
| 55 | Continental-scale conservation prioritization of African dragonflies | (Simaika et al., 2013) | Biological Conservation | 1 | Ensemble | No | Zonation | No |
| 56 | Accommodating Species Climate-Forced Dispersal and Uncertainties in Spatial Conservation Planning | (Lemes and Loyola, 2013) | Plos One | 6 | Ensemble | Yes | Zonation | Yes |
| 57 | Predictive habitat modelling of reef fishes with contrasting trophic ecologies | (Schmiing et al., 2013) | MEPS | 1 | - | No | Notation | No |
| 58 | Areas of Climate Stability of Species Ranges in the Brazilian Cerrado: Disentangling Uncertainties Through Time | (Terribile et al., 2012) | Natureza & Conservacao | 1 | Ensemble | No | Zonation | No |

| | | | | | | | | |
|---|---|---|---|---|---|---|---|---|
| 59 | Coexistence of mesopredators in an intact polar ocean ecosystem: The basis for defining a Ross Sea marine protected area | (Ballard et al., 2012) | Biological Conservation | 1 | - | No | Marxan | No |
| 60 | Ecological coherence of marine protected area networks: a spatial assessment using species distribution models | (Sundblad et al., 2011) | JOURNAL OF APPLIED ECOLOGY | 1 | - | Yes | Zonation | Yes |
| 61 | The importance of land use/land cover data in fish and mussel conservation planning | (Hopkins and Whiles, 2011) | ANNALES DE LIMNOLOGIE-INTERNATIONAL JOURNAL OF LIMNOLOGY | 1 | Best | No | Notation | No |
| 62 | Assessing conservation priorities of xenarthrans in Argentina | (Tognelli et al., 2011) | BIODIVERSITY AND CONSERVATION | 1 | Ensemble | No | Zonation | No |
| 63 | Lizards as conservation targets in Argentinean Patagonia | (Corbalán et al., 2011) | JOURNAL FOR NATURE CONSERVATION | 1 | Ensemble | No | Zonation | No |
| 64 | Simulating the effects of using different types of species distribution data in reserve selection | (Carvalho et al., 2010) | BIOLOGICAL CONSERVATION | 1 | - | No | Zonation | Yes |
| 65 | Optimizing resiliency of reserve networks to climate change: multispecies conservation planning in the Pacific | (Carroll et al., 2010) | GLOBAL CHANGE BIOLOGY | 1 | Best | Yes | Zonation | No |

| | | | | | | | | |
|---|---|---|---|---|---|---|---|---|
| | Northwest, USA | | | | | | | |
| 66 | Comparing alternative systematic conservation planning strategies against a politically driven conservation plan | (Meynard et al., 2009) | BIODIVERSITY AND CONSERVATIO N | 1 | Best | No | Marxan | No |
| 67 | Identifying important areas for butterfly conservation in Italy | (Girardello et al., 2009) | ANIMAL CONSERVATIO N | 1 | - | No | Zonation | No |
| 68 | Novel methods for the design and evaluation of marine protected areas in offshore waters | (Leathwick et al., 2008) | Conservation letters | 1 | Ensemble | No | Zonation | No |
| 69 | Identification of biodiversity conservation priorities using predictive modeling: An application for the equatorial pacific region of South America | (Peralvo et al., 2007) | BIODIVERSITY AND CONSERVATIO N | 1 | Best | No | SPOT | No |
| 70 | Conservation assessment and prioritization of areas in Northeast India: Priorities for amphibians and reptiles | (Pawar et al., 2007) | BIOLOGICAL CONSERVATIO N ; Conf??? | 1 | - | No | Resnet | No |
| 71 | Using landscape suitability models to reconcile conservation planning for two key forest predators | (Zielinski et al., 2006) | BIOLOGICAL CONSERVATIO N | 1 | - | No | Marxan | No |
| 72 | Connectivity, probabilities and persistence: Comparing reserve selection strategies | (van Teeffelen et al., 2006) | BIODIVERSITY AND CONSERVATIO N | 4 | - | No | Greedy | Yes |

| | | | | | | | | |
|---|---|---|---|---|---|---|---|---|
| 73 | Planning for climate change: Identifying minimum-dispersal corridors for the Cape proteaceae | (Williams et al., 2005) | CONSERVATION BIOLOGY | 1 | - | Yes | Greedy | No |
| 74 | Would climate change drive species out of reserves? An assessment of existing reserve-selection methods | (Araújo et al., 2004) | GLOBAL CHANGE BIOLOGY | 1 | - | Yes | Worldmap | No |
| 75 | Avoiding pitfalls of using SDMs in conservation planning | (Loiselle et al., 2003) | CONSERVATION BIOLOGY | 3 | - | No | Worldmap | Yes |

### Appendix 2

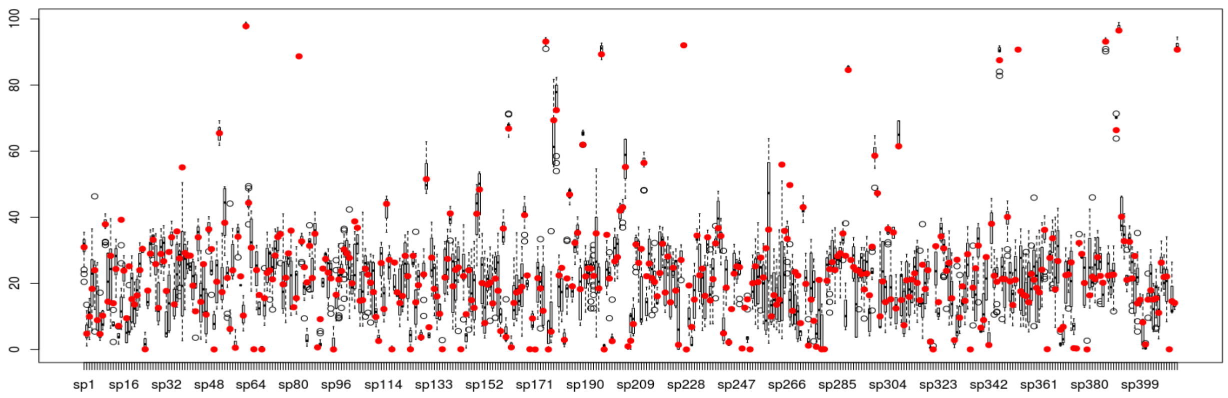


**Figure A2.1 :** Percentage of predicted species range across distribution models with good predictive performance. The points represent the predicted range from the ensemble model.

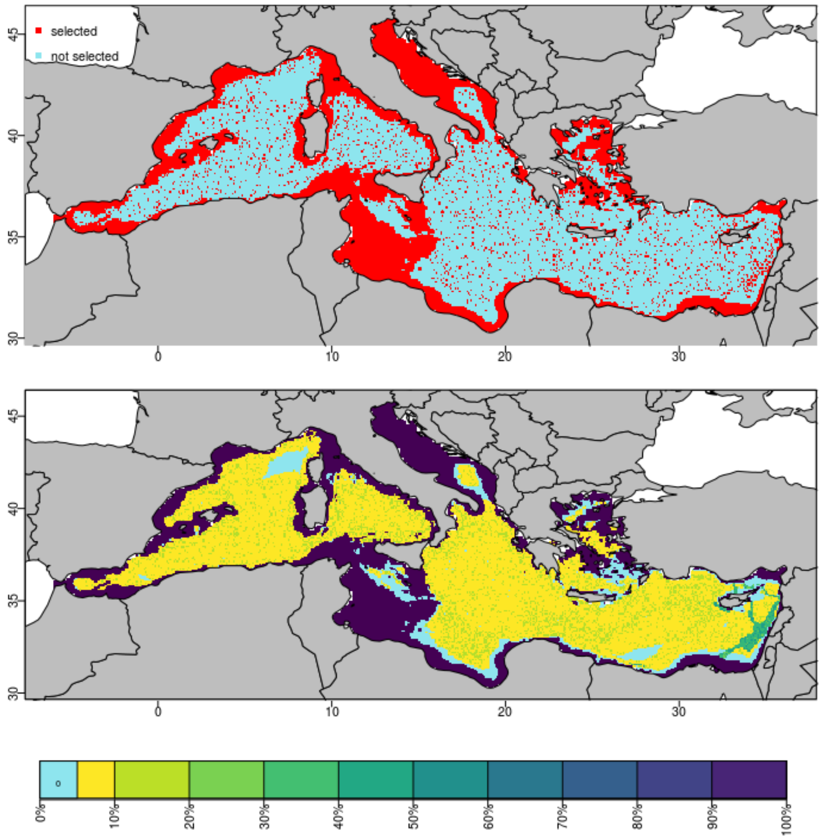


**Figure A2.2 :** Conservation outcomes following the pre-selection approach : (a) optimal solution from exact optimization (b) Priority ranking values across sub-optimal solutions

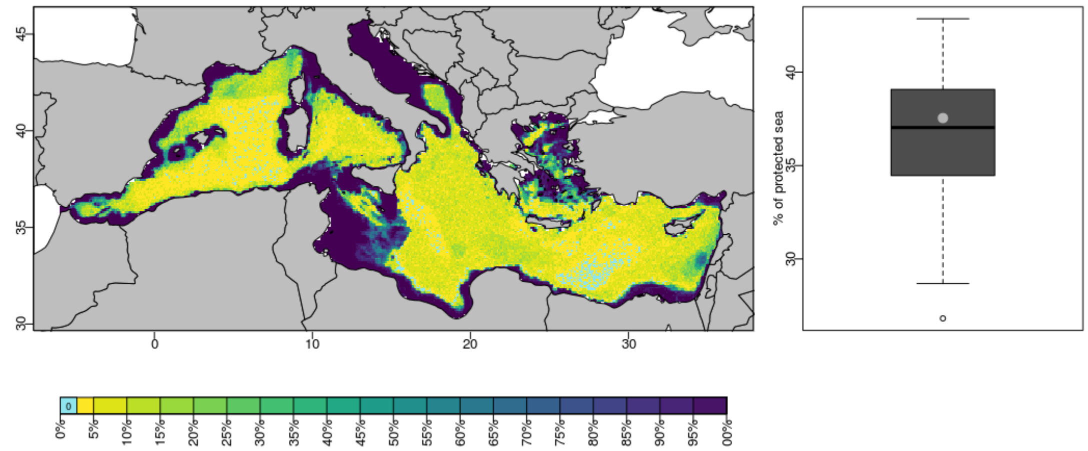


**Figure A2.3 :** Conservation outcomes following the post-selection approach : (a) Priority ranking values across optimal solutions build for each "distribution scenario". (b) Percentage of protected areas across those optimal solutions. The grey point represents the percentage of protected areas necessary to achieve species target based on ensemble model.

***Refrences***


Adams-Hosking, C., McAlpine, C.A., Rhodes, J.R., Moss, P.T., Grantham, H.S., 2015. Prioritizing Regions to Conserve a Specialist Folivore: Considering Probability of Occurrence, Food Resources, and Climate Change: Zonation prioritization for koala. Conservation Letters 8, 162–170. https://doi.org/10.1111/conl.12125

Alagador, D., Cerdeira, J.O., Ara?jo, M.B., 2016. Climate change, species range shifts and dispersal corridors: an evaluation of spatial conservation models. Methods in Ecology and Evolution 7, 853–866. https://doi.org/10.1111/2041-210X.12524

Alagador, D., Cerdeira, J.O., Ara?jo, M.B., 2014. Shifting protected areas: scheduling spatial priorities under climate change. Journal of Applied Ecology 51, 703–713. https://doi.org/10.1111/1365-2664.12230

Amorim, F., Carvalho, S.B., Honrado, J., Rebelo, H., 2014. Designing Optimized Multi-Species Monitoring Networks to Detect Range Shifts Driven by Climate Change: A Case Study with Bats in the North of Portugal. PLoS ONE 9, e87291. https://doi.org/10.1371/journal.pone.0087291

Araújo, M.B., Cabeza, M., Thuiller, W., Hannah, L., Williams, P.H., 2004. Would climate change drive species out of reserves? An assessment of existing reserve-selection methods. Global change biology 10, 1618–1626.

Ballard, G., Jongsomjit, D., Veloz, S.D., Ainley, D.G., 2012. Coexistence of mesopredators in an intact polar ocean ecosystem: The basis for defining a Ross Sea marine protected area. Biological Conservation 156, 72–82. https://doi.org/10.1016/j.biocon.2011.11.017

Beaudry, F., Ferris, M.C., Pidgeon, A.M., Radeloff, V.C., 2016. Identifying areas of optimal multispecies conservation value by accounting for incompatibilities between species. Ecological Modelling 332, 74–82. https://doi.org/10.1016/j.ecolmodel.2016.04.007

Bino, G., Ramp, D., Kingsford, R.T., 2014. Identifying minimal sets of survey techniques for multi-species monitoring across landscapes: an approach utilising species distribution models. International Journal of Geographical Information Science 28, 1674–1708. https://doi.org/10.1080/13658816.2013.871016

Bino, G., Ramp, D., Kingsford, R.T., 2013. Improving bioregional frameworks for conservation by including mammal distributions: IMPROVING BIOREGIONS BY USING MAMMAL ASSEMBLAGES. Austral Ecology 38, 393–404. https://doi.org/10.1111/j.1442-9993.2012.02423.x

Bond, N.R., Thomson, J.R., Reich, P., 2014. Incorporating climate change in conservation planning for freshwater fishes. Diversity and Distributions 20, 931–942. https://doi.org/10.1111/ddi.12213

Bush, A., Hermoso, V., Linke, S., Nipperess, D., Turak, E., Hughes, L., 2014. Freshwater conservation planning under climate change: demonstrating proactive approaches for Australian Odonata. Journal of Applied Ecology 51, 1273–1281. https://doi.org/10.1111/1365-2664.12295

Cadima, X., van Zonneveld, M., Scheldeman, X., Castañeda, N., Patiño, F., Beltran, M., Van Damme, P., 2014. Endemic wild potato (Solanum spp.) biodiversity status in Bolivia: Reasons for conservation concerns. Journal for Nature Conservation 22, 113–131. https://doi.org/10.1016/j.jnc.2013.09.007

Carroll, C., Dunk, J.R., Moilanen, A., 2010. Optimizing resiliency of reserve networks to climate change: multispecies conservation planning in the Pacific Northwest, USA. Global Change Biology 16, 891–904. https://doi.org/10.1111/j.1365-2486.2009.01965.x

Carvalho, S.B., Brito, J.C., Pressey, R.L., Crespo, E., Possingham, H.P., 2010. Simulating the effects of using different types of species distribution data in reserve selection. Biological Conservation 143, 426–438. https://doi.org/10.1016/j.biocon.2009.11.010

Carvalho, S.B., Gonçalves, J., Guisan, A., Honrado, J.P., 2016. Systematic site selection for multispecies monitoring networks. Journal of Applied Ecology 53, 1305–1316. https://doi.org/10.1111/1365-2664.12505

Chen, Y., Zhang, J., Jiang, J., Nielsen, S.E., He, F., 2017. Assessing the effectiveness of China's protected areas to conserve current and future amphibian diversity. Diversity and Distributions 23, 146–157. https://doi.org/10.1111/ddi.12508

Corbalán, V., Tognelli, M.F., Scolaro, J.A., Roig-Juñent, S.A., 2011. Lizards as conservation targets in Argentinean Patagonia. Journal for Nature Conservation 19, 60–67. https://doi.org/10.1016/j.jnc.2010.05.004

Davies, H.N., Beckley, L.E., Kobryn, H.T., Lombard, A.T., Radford, B., Heyward, A., 2016. Integrating Climate Change Resilience Features into the Incremental Refinement of an Existing Marine Park. PLOS ONE 11, e0161094. https://doi.org/10.1371/journal.pone.0161094

de Albuquerque, F.S., Beier, P., 2016. Downscaling patterns of complementarity to a finer resolution and its implications for conservation prioritization. Ecology and Evolution 6, 4032–4040. https://doi.org/10.1002/ece3.2190

de Albuquerque, F.S., Benito, B., Beier, P., Assunção-Albuquerque, M.J., Cayuela, L., 2015. Supporting underrepresented forests in Mesoamerica. Natureza & Conservação 13, 152–158. https://doi.org/10.1016/j.ncon.2015.02.001

Di Minin, E., Soutullo, A., Bartesaghi, L., Rios, M., Szephegyi, M.N., Moilanen, A., 2017. Integrating biodiversity, ecosystem services and socio-economic data to identify priority areas and landowners for conservation actions at the national scale. Biological Conservation 206, 56–64. https://doi.org/10.1016/j.biocon.2016.11.037

Dunn, J.C., Buchanan, G.M., Stein, R.W., Whittingham, M.J., McGowan, P.J.K., 2016. Optimising different types of biodiversity coverage of protected areas with a case study using Himalayan Galliformes. Biological Conservation 196, 22–30. https://doi.org/10.1016/j.biocon.2016.01.015

Fajardo, J., Lessmann, J., Bonaccorso, E., Devenish, C., Muñoz, J., 2014. Combined Use of Systematic Conservation Planning, Species Distribution Modelling, and Connectivity Analysis Reveals Severe Conservation Gaps in a Megadiverse Country (Peru). PLoS ONE 9, e114367. https://doi.org/10.1371/journal.pone.0114367

Faleiro, F.V., Loyola, R.D., 2013. Socioeconomic and political trade-offs in biodiversity conservation: a case study of the Cerrado Biodiversity Hotspot, Brazil. Diversity and Distributions 19, 977–987. https://doi.org/10.1111/ddi.12072

Fleishman, E., Thomson, J.R., Kalies, E.L., Dickson, B.G., Dobkin, D.S., Leu, M., 2014. Projecting current and future location, quality, and connectivity of habitat for breeding birds in the Great Basin. Ecosphere 5, art82. https://doi.org/10.1890/ES13-00387.1

Girardello, M., Griggio, M., Whittingham, M.J., Rushton, S.P., 2009. Identifying important areas for butterfly conservation ï¿½in Italy. Animal Conservation 12, 20–28. https://doi.org/10.1111/j.1469-1795.2008.00216.x

Gomez, C., Lawson, J., Kouwenberg, A., Moors-Murphy, H., Buren, A., Fuentes-Yaco, C., Marotte, E., Wiersma, Y., Wimmer, T., 2017. Predicted distribution of whales at risk: identifying priority areas to enhance cetacean monitoring in the Northwest Atlantic Ocean. Endangered Species Research 32, 437–458. https://doi.org/10.3354/esr00823

Hermoso, V., Kennard, M.J., Schmidt, D.J., Bond, N., Huey, J.A., Mondol, R.K., Jamandre, B.W., Hughes, J.M., 2016. Species distributions represent intraspecific genetic diversity of freshwater fish in conservation assessments. Freshwater Biology 61, 1707–1719. https://doi.org/10.1111/fwb.12810

Hopkins, R.L., Whiles, M.R., 2011. The importance of land use/land cover data in fish and mussel conservation planning. Annales de Limnologie - International Journal of Limnology 47, 199–209. https://doi.org/10.1051/limn/2011026

Huang, X., Li, F., Chen, J., 2016. Reserve network planning for fishes in the middle and lower Yangtze River basin by systematic conservation approaches. Science China Life Sciences 59, 312–324. https://doi.org/10.1007/s11427-015-4950-0

Lacasella, F., Marta, S., Singh, A., Stack Whitney, K., Hamilton, K., Townsend, P., Kucharik, C.J.,

Meehan, T.D., Gratton, C., 2017. From pest data to abundance-based risk maps combining eco-physiological knowledge, weather, and habitat variability. Ecological Applications 27, 575–588. https://doi.org/10.1002/eap.1467
Laze, K., Gordon, A., 2016. Incorporating natural and human factors in habitat modelling and spatial prioritisation for the <i>Lynx lynx martinoi</i> Web Ecology 16, 17–31. https://doi.org/10.5194/we-16-17-2016
Leach, K., Zalat, S., Gilbert, F., 2013. Egypt's Protected Area network under future climate change. Biological Conservation 159, 490–500. https://doi.org/10.1016/j.biocon.2012.11.025
Leathwick, J., Moilanen, A., Francis, M., Elith, J., Taylor, P., Julian, K., Hastie, T., Duffy, C., 2008. Novel methods for the design and evaluation of marine protected areas in offshore waters: Designing offshore MPAs. Conservation Letters 1, 91–102. https://doi.org/10.1111/j.1755-263X.2008.00012.x
Lemes, P., Loyola, R.D., 2013. Accommodating Species Climate-Forced Dispersal and Uncertainties in Spatial Conservation Planning. PLoS ONE 8, e54323. https://doi.org/10.1371/journal.pone.0054323
Lentini, P.E., Wintle, B.A., 2015. Spatial conservation priorities are highly sensitive to choice of biodiversity surrogates and species distribution model type. Ecography 38, 1101–1111. https://doi.org/10.1111/ecog.01252
Lessmann, J., Fajardo, J., Muñoz, J., Bonaccorso, E., 2016. Large expansion of oil industry in the Ecuadorian Amazon: biodiversity vulnerability and conservation alternatives. Ecology and Evolution 6, 4997–5012. https://doi.org/10.1002/ece3.2099
Lessmann, J., Muñoz, J., Bonaccorso, E., 2014. Maximizing species conservation in continental Ecuador: a case of systematic conservation planning for biodiverse regions. Ecology and Evolution 4, 2410–2422. https://doi.org/10.1002/ece3.1102
Loiselle, B.A., Howell, C.A., Graham, C.H., Goerck, J.M., Brooks, T., Smith, K.G., Williams, P.H., 2003. Avoiding Pitfalls of Using Species Distribution Models in Conservation Planning. Conservation Biology 17, 1591–1600. https://doi.org/10.1111/j.1523-1739.2003.00233.x
Long, A.L., Kettenring, K.M., Toth, R., 2017. Prioritizing Management of the Invasive Grass Common Reed ( *Phragmites* australis ) in Great Salt Lake Wetlands. Invasive Plant Science and Management 10, 155–165. https://doi.org/10.1017/inp.2017.20
Maleki, S., Soffianian, A.R., Soltani Koupaei, S., Saatchi, S., Pourmanafi, S., Sheikholeslam, F., 2016. Habitat mapping as a tool for water birds conservation planning in an arid zone wetland: The case study Hamun wetland. Ecological Engineering 95, 594–603. https://doi.org/10.1016/j.ecoleng.2016.06.115
Mateo, R.G., de la Estrella, M., Felicísimo, Á.M., Muñoz, J., Guisan, A., 2013. A new spin on a compositionalist predictive modelling framework for conservation planning: A tropical case study in Ecuador. Biological Conservation 160, 150–161. https://doi.org/10.1016/j.biocon.2013.01.014
McGowan, J., Hines, E., Elliott, M., Howar, J., Dransfield, A., Nur, N., Jahncke, J., 2013. Using Seabird Habitat Modeling to Inform Marine Spatial Planning in Central California's National Marine Sanctuaries. PLoS ONE 8, e71406. https://doi.org/10.1371/journal.pone.0071406
Mendoza, E., Fuller, T.L., Thomassen, H.A., Buermann, W., Ramírez-Mejía, D., Smith, T.B., 2013. A preliminary assessment of the effectiveness of the Mesoamerican Biological Corridor for protecting potential Baird's tapir ( *Tapirus* bairdii ) habitat in southern Mexico. Integrative Zoology 8, 35–47. https://doi.org/10.1111/1749-4877.12005
Meynard, C.N., Howell, C.A., Quinn, J.F., 2009. Comparing alternative systematic conservation planning strategies against a politically driven conservation plan. Biodiversity and Conservation 18, 3061–3083. https://doi.org/10.1007/s10531-009-9625-3
Morán-Ordóñez, A., Whitehead, A.L., Luck, G.W., Cook, G.D., Maggini, R., Fitzsimons, J.A., Wintle, B.A., 2017. Analysis of Trade-Offs Between Biodiversity, Carbon Farming and Agricultural Development in Northern Australia Reveals the Benefits of Strategic Planning:

Spatial planning northern Australian Savannas. Conservation Letters 10, 94–104. https://doi.org/10.1111/conl.12255
Nieto, C., Ovando, X.M.C., Loyola, R., Izquierdo, A., Romero, F., Molineri, C., Rodríguez, J., Rueda Martín, P., Fernández, H., Manzo, V., Miranda, M.J., 2017. The role of macroinvertebrates for conservation of freshwater systems. Ecology and Evolution 7, 5502–5513. https://doi.org/10.1002/ece3.3101
Passoni, G., Rowcliffe, J.M., Whiteman, A., Huber, D., Kusak, J., 2017. Framework for strategic wind farm site prioritisation based on modelled wolf reproduction habitat in Croatia. European Journal of Wildlife Research 63. https://doi.org/10.1007/s10344-017-1092-7
Pawar, S., Koo, M.S., Kelley, C., Ahmed, M.F., Chaudhuri, S., Sarkar, S., 2007. Conservation assessment and prioritization of areas in Northeast India: Priorities for amphibians and reptiles. Biological Conservation 136, 346–361. https://doi.org/10.1016/j.biocon.2006.12.012
Peralvo, M., Sierra, R., Young, K.R., Ulloa, C.U.-, 2007. Identification of Biodiversity Conservation Priorities using Predictive Modeling: An Application for the Equatorial Pacific Region of South America. Biodiversity and Conservation 16, 2649–2675. https://doi.org/10.1007/s10531-006-9077-y
Pollock, L.J., Rosauer, D.F., Thornhill, A.H., Kujala, H., Crisp, M.D., Miller, J.T., McCarthy, M.A., 2015. Phylogenetic diversity meets conservation policy: small areas are key to preserving eucalypt lineages. Philosophical Transactions of the Royal Society B: Biological Sciences 370, 20140007–20140007. https://doi.org/10.1098/rstb.2014.0007
Reside, A.E., VanDerWal, J., Moran, C., 2017. Trade-offs in carbon storage and biodiversity conservation under climate change reveal risk to endemic species. Biological Conservation 207, 9–16. https://doi.org/10.1016/j.biocon.2017.01.004
Runge, C.A., Tulloch, A.I.T., Possingham, H.P., Tulloch, V.J.D., Fuller, R.A., 2016. Incorporating dynamic distributions into spatial prioritization. Diversity and Distributions 22, 332–343. https://doi.org/10.1111/ddi.12395
Schmiing, M., Afonso, P., Tempera, F., Santos, R., 2013. Predictive habitat modelling of reef fishes with contrasting trophic ecologies. Marine Ecology Progress Series 474, 201–216. https://doi.org/10.3354/meps10099
Simaika, J.P., Samways, M.J., Kipping, J., Suhling, F., Dijkstra, K.-D.B., Clausnitzer, V., Boudot, J.-P., Domisch, S., 2013. Continental-scale conservation prioritization of African dragonflies. Biological Conservation 157, 245–254. https://doi.org/10.1016/j.biocon.2012.08.039
Smith, A., Schoeman, M.C., Keith, M., Erasmus, B.F.N., Monadjem, A., Moilanen, A., Di Minin, E., 2016. Synergistic effects of climate and land-use change on representation of African bats in priority conservation areas. Ecological Indicators 69, 276–283. https://doi.org/10.1016/j.ecolind.2016.04.039
Stralberg, D., Bayne, E.M., Cumming, S.G., Sólymos, P., Song, S.J., Schmiegelow, F.K.A., 2015. Conservation of future boreal forest bird communities considering lags in vegetation response to climate change: a modified refugia approach. Diversity and Distributions 21, 1112–1128. https://doi.org/10.1111/ddi.12356
Sundblad, G., Bergström, U., Sandström, A., 2011. Ecological coherence of marine protected area networks: a spatial assessment using species distribution models: Assessing eco-coherence of MPA networks. Journal of Applied Ecology 48, 112–120. https://doi.org/10.1111/j.1365-2664.2010.01892.x
Terribile, L.C. et al., 2012. Areas of climate stability of species ranges in the Brazilian Cerrado: disentangling uncertainties through time. Natureza & Conservação 10, 152–159. https://doi.org/10.4322/natcon.2012.025
Tognelli, M.F., Abba, A.M., Bender, J.B., Seitz, V.P., 2011. Assessing conservation priorities of xenarthrans in Argentina. Biodiversity and Conservation 20, 141–151. https://doi.org/10.1007/s10531-010-9951-5
van Teeffelen, A.J.A., Cabeza, M., Moilanen, A., 2006. Connectivity, Probabilities and Persistence:

Comparing Reserve Selection Strategies. Biodiversity & Conservation 15, 899–919. https://doi.org/10.1007/s10531-004-2933-8
Veloz, S.D., Nur, N., Salas, L., Jongsomjit, D., Wood, J., Stralberg, D., Ballard, G., 2013. Modeling climate change impacts on tidal marsh birds: Restoration and conservation planning in the face of uncertainty. Ecosphere 4, art49. https://doi.org/10.1890/ES12-00341.1
Walther, B.A., Pirsig, L.H., 2017. Determining conservation priority areas for Palearctic passerine migrant birds in sub-Saharan Africa. Avian Conservation and Ecology 12. https://doi.org/10.5751/ACE-00934-120102
Wan, J., Wang, C., Han, S., Yu, J., 2014a. Planning the priority protected areas of endangered orchid species in northeastern China. Biodiversity and Conservation 23, 1395–1409. https://doi.org/10.1007/s10531-014-0671-0
Wan, J., Wang, C., Yu, J., Nie, S., Han, S., Liu, J., Zu, Y., Wang, Q., 2016. Developing conservation strategies for Pinus koraiensis and Eleutherococcus senticosus by using model-based geographic distributions. Journal of Forestry Research 27, 389–400. https://doi.org/10.1007/s11676-015-0170-5
Wan, J., Wang, C., Yu, J., Nie, S., Han, S., Zu, Y., Chen, C., Yuan, S., Wang, Q., 2014b. Model-based conservation planning of the genetic diversity of *Phellodendron* amurense Rupr due to climate change. Ecology and Evolution 4, 2884–2900. https://doi.org/10.1002/ece3.1133
Wan, J.-Z., Wang, C.-J., Yu, F.-H., 2017. Spatial conservation prioritization for dominant tree species of Chinese forest communities under climate change. Climatic Change 144, 303–316. https://doi.org/10.1007/s10584-017-2044-7
Wang, C.-J., Wan, J.-Z., Zhang, Z.-X., Zhang, G.-M., 2016. Identifying appropriate protected areas for endangered fern species under climate change. SpringerPlus 5. https://doi.org/10.1186/s40064-016-2588-4
Whitehead, A.L., Kujala, H., Ives, C.D., Gordon, A., Lentini, P.E., Wintle, B.A., Nicholson, E., Raymond, C.M., 2014. Integrating Biological and Social Values When Prioritizing Places for Biodiversity Conservation: Conservation Planning with Social Data. Conservation Biology 28, 992–1003. https://doi.org/10.1111/cobi.12257
Whitehead, A.L., Kujala, H., Wintle, B.A., 2017. Dealing with Cumulative Biodiversity Impacts in Strategic Environmental Assessment: A New Frontier for Conservation Planning: Cumulative biodiversity impact assessments. Conservation Letters 10, 195–204. https://doi.org/10.1111/conl.12260
Williams, J.W., Kharouba, H.M., Veloz, S., Vellend, M., McLachlan, J., Liu, Z., Otto-Bliesner, B., He, F., 2013. The ice age ecologist: testing methods for reserve prioritization during the last global warming: Reserve selection and the ice age ecologist. Global Ecology and Biogeography 22, 289–301. https://doi.org/10.1111/j.1466-8238.2012.00760.x
Williams, P., Hannah, L., Andelman, S., Midgley, G., Araújo, M., Hughes, G., Manne, L., Martinez-Meyer, E., Pearson, R., 2005. Planning for Climate Change: Identifying Minimum-Dispersal Corridors for the Cape Proteaceae: *Planning* for Climate Change. Conservation Biology 19, 1063–1074. https://doi.org/10.1111/j.1523-1739.2005.00080.x
Winiarski, K.J., Miller, D.L., Paton, P.W.C., McWilliams, S.R., 2014. A spatial conservation prioritization approach for protecting marine birds given proposed offshore wind energy development. Biological Conservation 169, 79–88. https://doi.org/10.1016/j.biocon.2013.11.004
Zielinski, W.J., Carroll, C., Dunk, J.R., 2006. Using landscape suitability models to reconcile conservation planning for two key forest predators. Biological Conservation 133, 409–430. https://doi.org/10.1016/j.biocon.2006.07.003
Zwiener, V.P., Padial, A.A., Marques, M.C.M., Faleiro, F.V., Loyola, R., Peterson, A.T., 2017. Planning for conservation and restoration under climate and land use change in the Brazilian Atlantic Forest. Diversity and Distributions 23, 955–966. https://doi.org/10.1111/ddi.12588
